# Gut Bacterial Dysbiosis and Instability is Associated with the Onset of Complications and Mortality in COVID-19

**DOI:** 10.1101/2021.10.08.463613

**Authors:** David Schult, Sandra Reitmeier, Plamena Koyumdzhieva, Tobias Lahmer, Moritz Middelhoff, Johanna Erber, Jochen Schneider, Juliane Kager, Marina Frolova, Julia Horstmann, Lisa Fricke, Katja Steiger, Moritz Jesinghaus, Klaus-Peter Janssen, Ulrike Protzer, Klaus Neuhaus, Roland M. Schmid, Dirk Haller, Michael Quante

## Abstract

**Objective:** There is a growing debate about the involvement of the gut microbiome in COVID-19, although it is not conclusively understood whether the microbiome has an impact on COVID-19, or vice versa, especially as analysis of amplicon data in hospitalized patients requires sophisticated cohort recruitment and integration of clinical parameters. Here, we analyzed fecal and saliva samples from SARS-CoV-2 infected and post COVID-19 patients and controls considering multiple influencing factors during hospitalization.

**Design:** 16S rRNA gene sequencing was performed on fecal and saliva samples from 108 COVID-19 and 22 post COVID-19 patients, 20 pneumonia controls and 26 asymptomatic controls. Patients were recruited over the first and second corona wave in Germany and detailed clinical parameters were considered. Serial samples per individual allowed intra-individual analysis.

**Results:** We found the gut and oral microbiota to be altered depending on number and type of COVID-19-associated complications and disease severity. The occurrence of individual complications was correlated with low-risk (e.g., *Faecalibacterium prausznitzii*) and high-risk bacteria (e.g., *Parabacteroides*). We demonstrated that a stable gut bacterial composition was associated with a favorable disease progression. Based on gut microbial profiles, we identified a model to estimate mortality in COVID-19.

**Conclusion:** Gut microbiota are associated with the occurrence of complications in COVID-19 and may thereby influencing disease severity. A stable gut microbial composition may contribute to a favorable disease progression and using bacterial signatures to estimate mortality could contribute to diagnostic approaches. Importantly, we highlight challenges in the analysis of microbial data in the context of hospitalization.

## Introduction

The global pandemic caused by the new severe acute respiratory syndrome coronavirus 2 (SARS-CoV-2) brought the health systems to its limitations. The disease is characterized by respiratory symptoms although there is increasing evidence of gastrointestinal (GI) tract involvement[1, 2, 3]. At 15-39%, nausea, vomiting, and diarrhea are relatively common in COVID-19[4] and a proportion of patients reports only gastrointestinal symptoms[5]. The virus itself is not limited to the lungs but replicates in human enterocytes[6] and is detectable in the patients’ fecal samples[1, 7]. GI symptoms in patients with COVID-19 appear to be associated with increased disease severity and complications[8], although the underlying causes are not understood. Recent studies suggest that an altered microbial composition correlates with COVID-19 disease severity and inflammatory response to the disease[9, 10].

Common complications of COVID-19 include venous thromboembolism[11, 12], hemodynamic instability[13], and acute kidney injury[14]. Particularly in severe cases, an excessive and prolonged immune response to the virus is thought to be a catalyst of severity[15, 16].

The composition of the gut microbiota plays a critical role in the immunological homeostasis of the human body[17, 18]. It is known that the microbiome of the human gut is sensitive to changes in the hosts’ environment[19]. In addition to antibiotic use, diet[20], and geographical differences[21, 22], critically ill patients show a rapid depletion of health-promoting organisms[23].

The study examined the impact of gut and oral microbiota on complication rate and outcome and, conversely, how hospitalization affects the gut microbial composition in this cohort.

## MATERIAL AND METHODS

### Study Cohorts

The study population consists of 4 groups: (1) 108 patients with laboratory confirmed SARS-CoV-2 infection, (2) 22 patients post COVID-19 who had cleared the virus and were tested negative at first sampling, (3) 20 symptomatic pneumonia controls (SC) and (4) 26 age and gender matched asymptomatic controls (AC) **(Table 1, Figure 1 A)**. Altogether, 251 stool samples and 160 saliva samples were examined. Serial samples were collected to investigate intra-individual changes over time. A total of 25 and 15 COVID-19 patients, 11 and 5 post COVID-19 patients and 3 and 2 SC provided serial stool and saliva samples, respectively **(Table 1)**. The SC patients were admitted with respiratory symptoms of community-acquired-pneumonia (CAP) and were tested negative for SARS-CoV-2. Patients in the AC group were considered SARS-CoV-2 negative as they presented mainly for screening colonoscopy and showed no symptoms of SARS-CoV-2 infection. To minimize potential influencing factors on the microbiota in the AC cohort, patients with active cancer, inflammatory bowel disease (IBD), oncologic therapy or antibiotic intake at the time of examination or within 6 months prior were excluded. Endoscopic examination and pathology reports from colon biopsies had to be unremarkable.

**Table 1.**
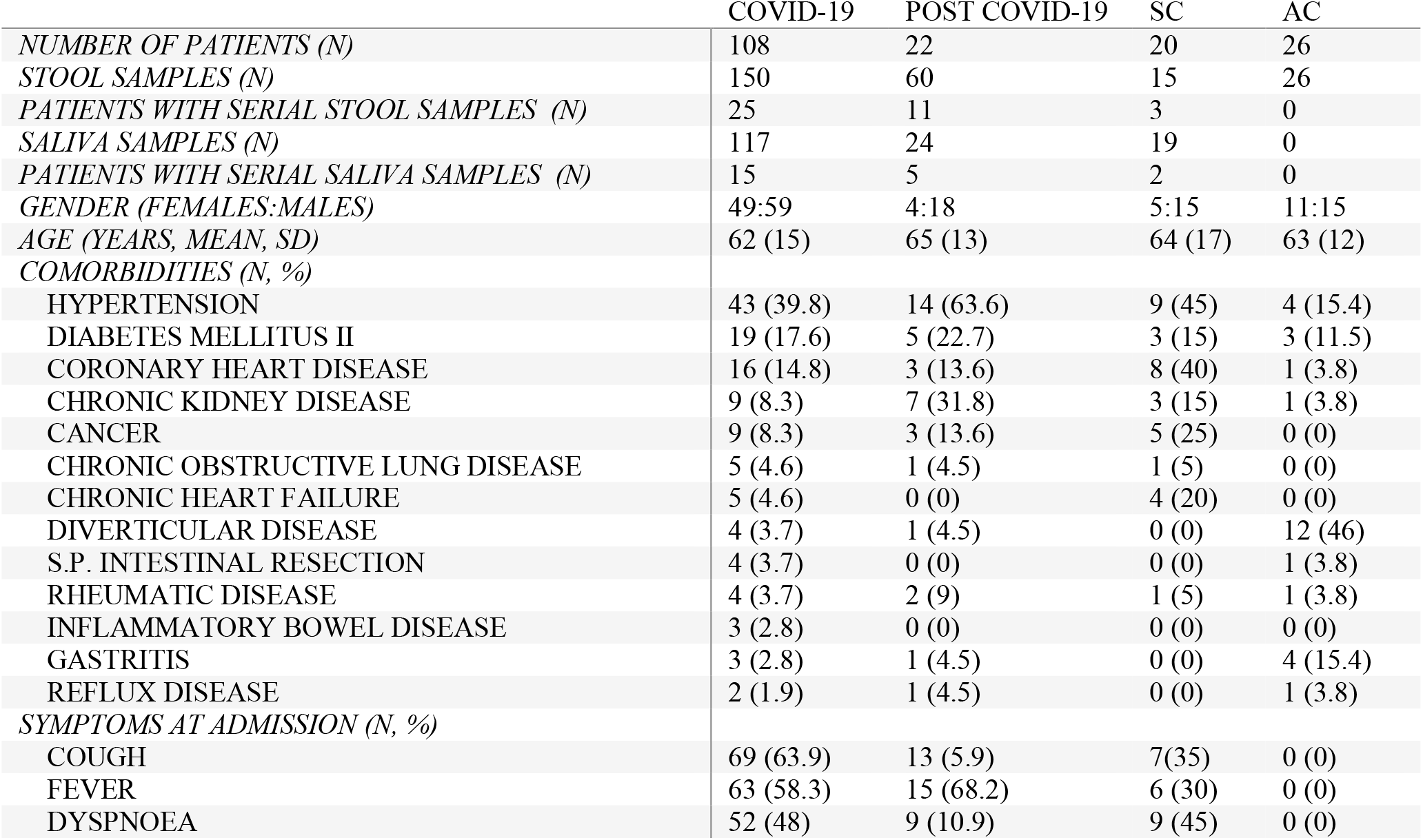
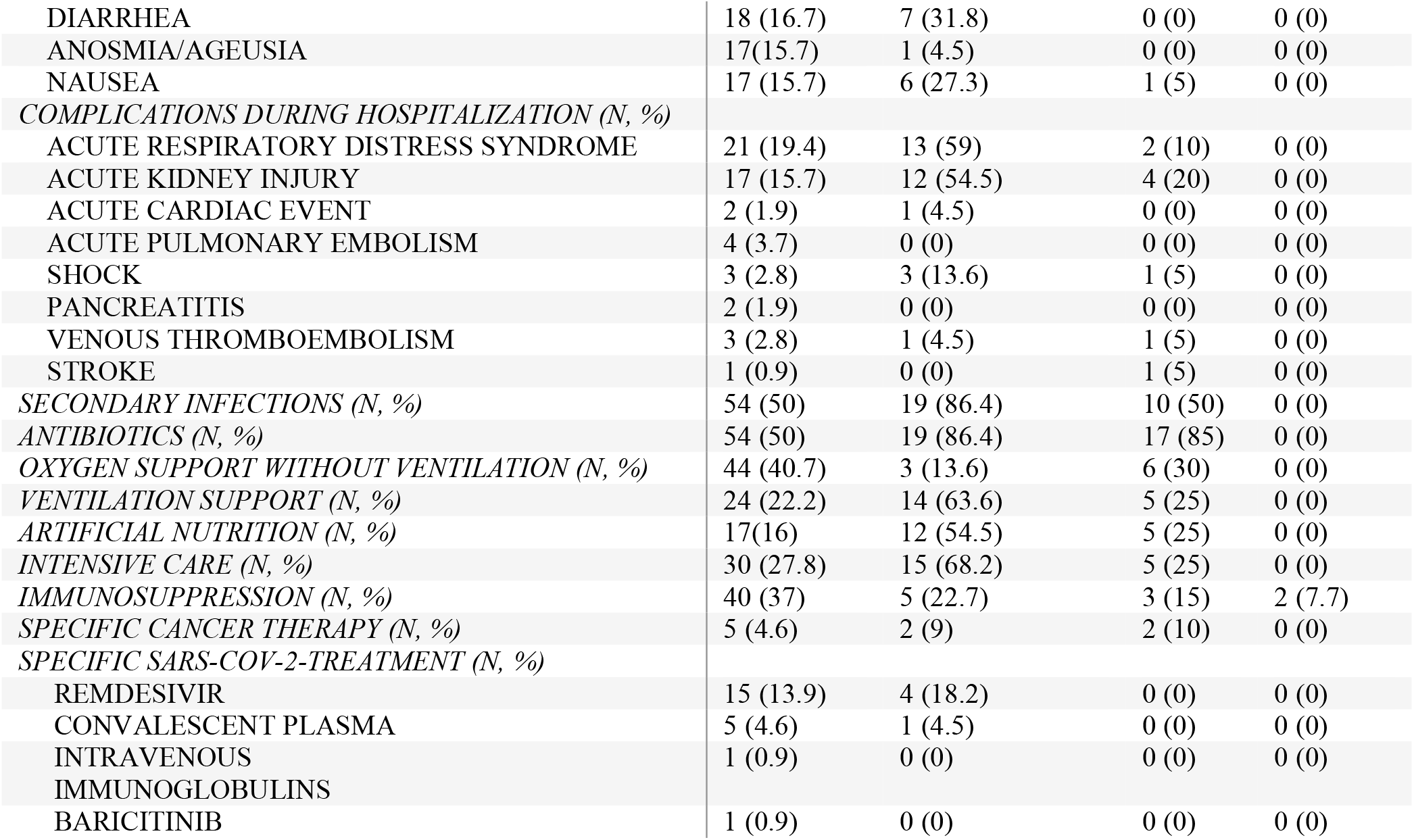
Demographic and Clinical Characteristics of the Study Population.

**Figure 1.**
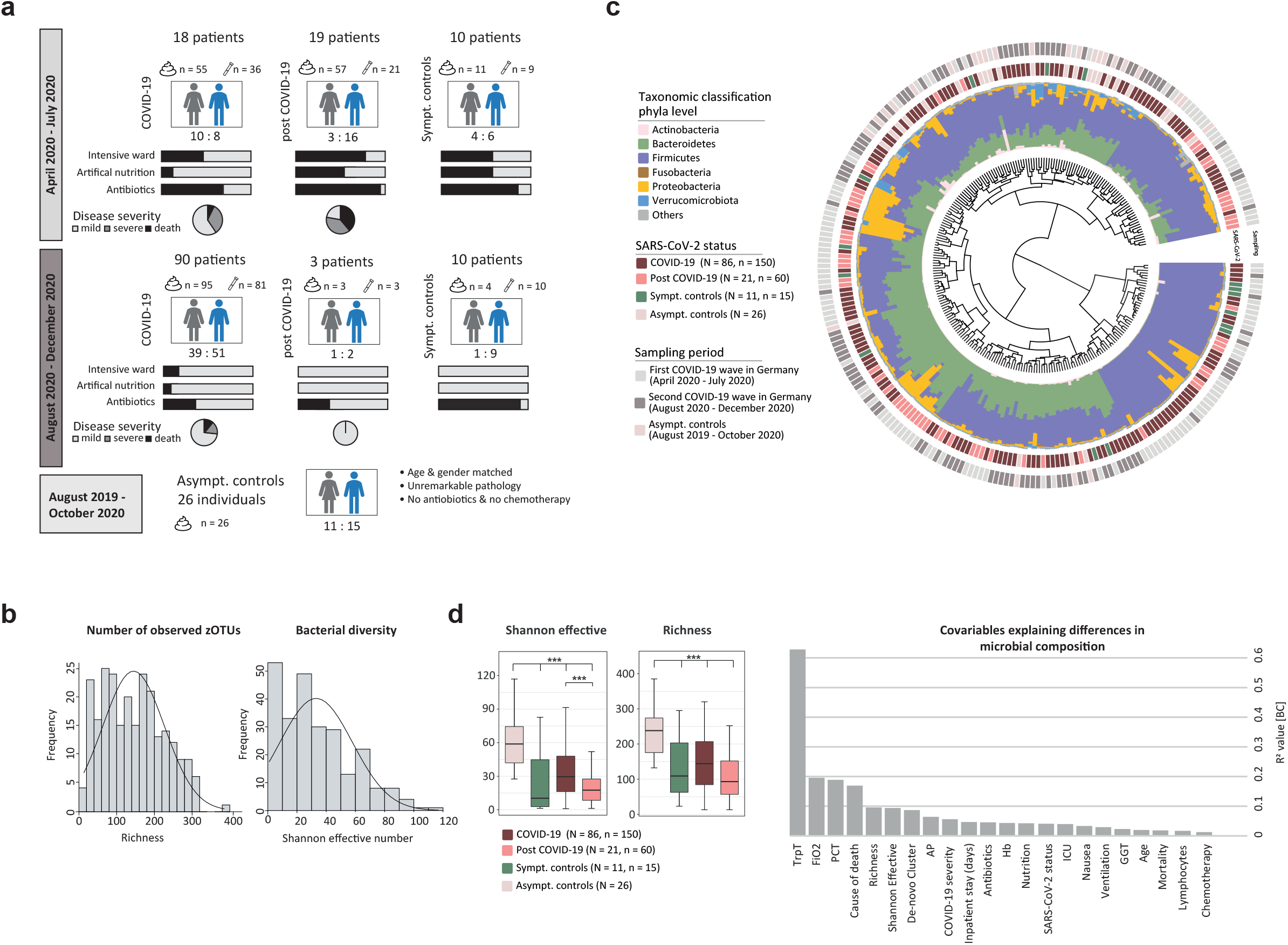
Microbial Composition of the Gut Observed in the Cohort. **A** Overview of study design. Stool and saliva samples are indicated. **B** *Alpha*-diversity of all samples of all patients. Left histogram shows richness and right histogram Shannon effective number of species. **C** Phylogenetic tree calculated by generalized Unifrac distances for all samples of all patients. Stacked barplots show taxonomic distribution on phyla level. Inner label shows SARS-CoV-2 status and outer label indicates the sampling time phase. **D** Left, *alpha*-diversity stratified according to SARS-CoV-2 status for all samples of all patients, showing Shannon effective numbers and richness. Right, barplots show effect modifiers significantly contributing to microbial diversity in all samples. Y-axis shows the R^2^ value calculated based on Bray-Curtis distance for COVID-19, post COVID-19 and SC.

### Patient Recruitment and Sampling

Acquisition of patients was conducted at the university hospital Klinikum rechts der Isar, Technical University Munich, Germany. COVID-19 patients, post COVID-19 patients and SC were prospectively recruited between April 2020 to July 2020 (first COVID-19 wave in Germany) and August 2020 to December 2020 (second COVID-19 wave in Germany), whereas the AC group was prospectively recruited between August 2019 and October 2020. Because these were control patients in an intestinal microbiome-only study, saliva was not obtained **(Figure 1 A)**. Stool, saliva and blood samples were collected at least once per week during the inpatient stay. To ensure follow-up and bio-sample collection after discharge, patients were invited to follow-up visits. SARS-CoV-2 infection was confirmed by quantitative reverse transcription PCR (RT-qPCR), performed on nasopharyngeal swabs. For the post COVID-19 patients the first stool sample was collected on average 30 days after the negative PCR. In the AC group, stool samples were collected either before or 6 weeks after bowel preparation for colonoscopy. To characterize the disease activity, laboratory parameters and data regarding fraction of inspired oxygen (FiO2), ventilation mode, diet, intensive or normal ward and antibiotic use were collected at each time point of stool or saliva sampling.

### Classifications

Patients with COVID-19 or post COVID-19 were classified into 3 groups based on the WHO ordinal scale for clinical improvement for hospitalized patients with COVID-19[24], which has been used in other COVID-19 studies[25]: (i) mild disease, composed of patients with no oxygen therapy (score 3) or oxygen by mask or nasal prongs (score 4); (ii) severe disease, including patients requiring non-invasive ventilation or high-flow oxygen (score 5), intubation and mechanical ventilation (score 6) or ventilation and additional organ support (score 7), and (iii) fatal disease (death, score 8). Ventilation mode during inpatient stay was divided in two groups: (i) Oxygen via nasal prongs, and (ii) mechanically ventilated either pressure controlled (PC) or pressure assisted (PA) and tracheostomy (TS) after long period of intubation. Considering the varying impact of different antibiotics on the gut microbiota, antibiotic therapy was classified according to their spectrum of activity **(Supplementary Table 1)**. Patients were either fed normally or with formulated food via gastric tube in combination with or without parenteral nutrition (summarized in tube feeding).

### Ethical Approval

All patients provided written informed consent. The study was conducted in accordance with the declaration of Helsinki and approved by the ethics committee of the Technical University Hospital of Munich (221/20 S-SR).

### Sample Preparation and 16S rRNA Gene Sequencing

Faecal and saliva samples were stored in a solution to stabilize DNA (MaGix PBI, Microbiomix GmbH, Regensburg Germany). Sample preparation and paired-end sequencing was performed on an Illumina MiSeq targeting the V3V4 region of the 16S rRNA gene. Detailed description of the methods is published[26]. Raw FASTQ files were processed using the NGSToolkit (https://github.com/TUM-Core-Facility-Microbiome/ngstoolkit) based on USEARCH 11[27] to generate denoised zero-radiation operational-taxonomic units (zOTUs).

### Statistical Analysis

Differences in relative abundance of taxa and/or zOTUs were determined by Kruskal-Wallis Rank Sum test for multiple groups and Wilcoxon Rank Sum test for pairwise comparison. Differences in prevalence were determined by a non-linear Fisher Exact test. Spearman correlation was calculated for associations and continuous variables.

Similarity between samples was estimated based on a distance matrix using generalized UniFrac. Significance between groups, effect modifier, and confounder were determined by a permutational multivariate analysis of variances (*adonis* function of the R-package vegan). For all analyses, p-values were corrected for multiple testing according to Benjamini-Hochberg correction.

The explained variation of co-variables was determined by calculating R^2^ values and were considered as significant with a p-value ≤0.05. A *random forest* model was used to classify binary outcome variables based on microbial composition with a 5-fold cross validation by using randomForest from the R package randomForest v4.6-14. To receive a robust and generalizable classification model, the machine-learning algorithm was applied 100-times iteratively. Based on out-of-bag error rates and Gini index, the most important features were selected for each iteration using *rfcv* from the R package randomForest v4.6-14. Features, which appeared in all 100 random forest models, were considered as classification features for the final model. A generalized linear model for binomial distribution and binary outcome (logit) was generated using the previously selected features.

## RESULTS

### Association of SARS-CoV-2 Status with the Gut Microbiota

Analysis of the gut microbiota was performed on 251 stool samples (n = 251) from 144 patients (N = 144), of which were 86 COVID-19 patients (n = 150 samples), 21 post COVID-19 patients (n = 60), 11 SC (n = 15) and 26 AC (n = 26) **(Figure 1 A)**. No bias due to sampling phases was observed allowing a combined analysis of the two COVID-19 waves.

Phylogenetic analysis of patient’s microbial profile showed no cluster according to SARS-CoV-2 status. Nevertheless, some patients were found to have an increased relative abundance of Proteobacteria which was mainly observed with COVID-19 and post COVID-19 samples **(Figure 1 C)**. The analysis of *alpha-*diversity revealed a not normally distributed number of observed species and bacterial diversity **(Figure 1 B)**. The number of observed species was reduced in active COVID-19 (richness 133 ± 90) and post COVID-19 patients (richness 103 ± 60) compared to AC (richness 219 ± 68), and bacterial diversity showed a reduced Shannon effective number in SC **(Figure 1 D)**.

Considering only the first sampling time point (T1) per individual revealed that parameters related to patient’s health were important effect modifiers **(Figure 1 D)**. Interestingly, even though the SARS-CoV-2 status alone did not show a clear pattern in the phylogenetic tree, the detection of SARS-CoV-2 in nasopharyngeal swabs significantly influenced the gut microbiota (R^2^ = 0.04, p = 0.001), as well as disease related variables, e.g. the disease severity (R^2^ = 0.05, p = 0.001).

### Evaluation of Confounding Factors

Although variables known to influence the microbial composition of the gut such as antibiotics or chemotherapy, appeared to be significant influencing factors **(Figure 1 D)**, none of the tested variables were confounders within the analysed cohort **(Supplementary Table 2)**. Particular attention was paid to variables related to hospitalization such as artificial feeding, critical care and antibiotic treatment. Since most patients were treated with different groups of antibiotics, we could not elucidate the influence of a specific antibiotic subgroup on the composition of the gut microbiota. Additionally, patients’ comorbidities and disease history was tested for confounding, considering type 2 diabetes[28, 29], inflammtory bowel disease[30], cancer, as well as chemotherapy and immunotherapy[31] within 6 months before stool sampling, or bowel resection[32] **(Supplementary Table 3)**. We further tested whether age and gender, specific SARS-CoV-2 treatment (remdesivir, convalescent plasma, intravenous immunoglobulins, or baricitinib), immunosuppressive therapy, or secondary infections introduced bias in the microbial analysis. Of note, critically ill patients with complications, compared to mild courses, were mainly treated at the ICU and received antibiotics **(Supplementary Table 4)**. However, within the cohort none of the above mentioned variables had a confounding effect in the analysis of the microbial composition related to COVID-19.

### Disease Severity and Progression Are Related to Altered Gut Microbiota

Disease severity according to WHO ordinal scale for clinical improvement significantly influences the gut bacterial composition of stool samples (p = 0.001) **(Figure 2 A)**. *Beta*-diversity clearly demonstrated a shift of bacterial profiles comparing controls with COVID-19 and post COVID-19 patients. Thereby, the bacterial composition of patients with a mild disease was more similar to SC and AC and a more severe disease showed a microbial composition more similar to patients who died due to COVID-19. A number of stool samples clustered independently in patients with severe and fatal COVID-19 disease, as well as a few mild courses and SC (**Figure 2 A**, left cluster). However, patients with mild disease in this cluster, or SC, showed no similarities for clinical or laboratory parameters with severe cases. None of the AC samples fell within this cluster.

**Figure 2.**
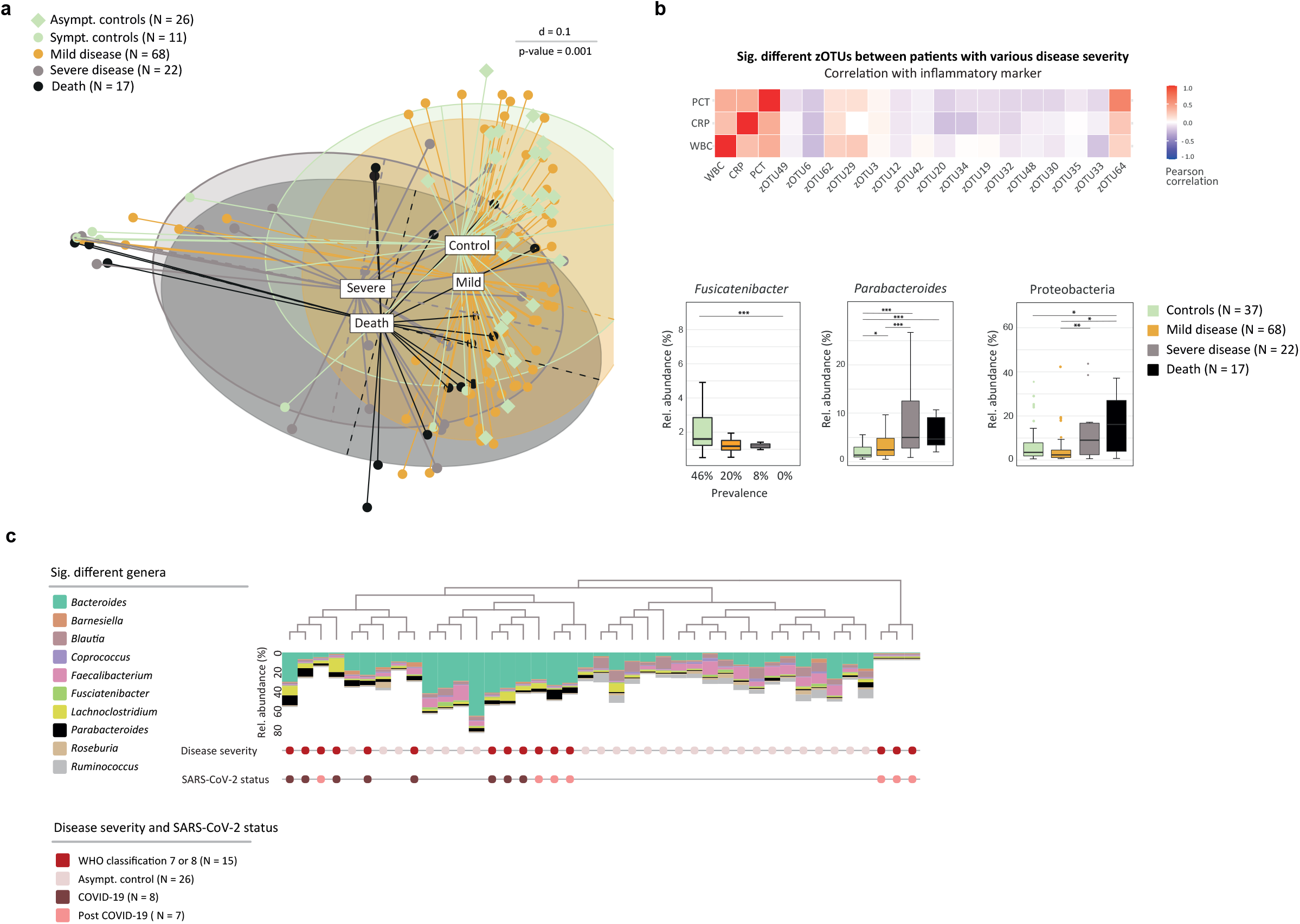
Microbial Profile of the Gut is Associated with Disease Severity. **A** MDS plot calculated on generalized UniFrac distance stratifying all patients (samples from the first time point, T1, only) according to disease severity. **B** Heatmap shows significant different taxa between COVID-19, post COVID-19 and SC patients with a different disease severity in correlation to inflammatory biomarkers. WBC, CRP and PCT. Boxplots show significantly different taxa according to disease severity. *Fusicatenibacter* shows differences in prevalence (p-value = 0.02), the genus *Parabacteroides* and phylum Protobacteria are significantly different in their relative abundance (p-value ≤ 0.001). **C** Dendrogram shows generalized UniFrac distances between a subset of COVID-19 and post COVID-19 patients, fulfilling certain criteria of a high inflammatory and severe disease, and AC for the sampling at T1. Stacked barplots display the relative abundance values of bacteria most significantly different.

Differentiation analysis revealed zOTUs **(Supplementary Table 5)**, which were significantly different between study groups and correlated with markers of inflammation, such as white blood cells counts (WBC), C-reactive protein (CRP) and procalcitonin (PCT) **(Figure 2 B)**. Here, *Clostridium innocuum* (zOTU62), *Ruthenibacterium lactatiformans* (zOTU29), and *Alistipes finegoldii* (zOTU64) correlated positively with inflammatory markers and continue to show a significantly increased relative abundance or prevalence in patients with a severe disease progression. Negatively correlated zOTUs were significantly decreased in severe and fatal cases of COVID-19 and post COVID-19, such as *Faecalibacterium prausnitzii* (zOTU20), *Blautia luti* (zOTU6), *Dorea longicatena* (zOTU32), *Gemmiger formicilis* (zOTU30), and *Alistipes putredinis* (zOTU33). In addition, *Fusicatenibacter* showed a significantly reduced prevalence in severe cases and was totally absent in patients who died **(Figure 2 B)**. On the other hand, *Parabacteroides* significantly increased with a more severe disease **(Figure 2 B)**. *Beta*-diversity already showed some accumulation of patients with an increased relative abundance of Protobacteria **(Figure 1 B)**, which was also found to be increased in severe COVID-19 cases **(Figure 2 B)**.

To analyse the associations of the gut microbial composition with COVID-19 severity in greater depth, we defined a subset of patients with certain criteria. This included patients presenting with high inflammatory parameters (CRP ≥ 10 mg/dl, PCT ≥ 5 ng/ml, WBC ≥ 15 G/l), FiO2 ≥ 40%, requiring mechanical ventilation (PC, PA, TS), were treated at the ICU, and had at least one complication. In addition, WHO disease severity was set to ≥6. Overall, 15 male patients met the criteria (COVID-19, N = 8; post COVID-19, N = 7) and all of them died, 13 due to acute respiratory distress syndrome (ARDS) and 2 of them due to cerebral haemorrhage. Stratification according to disease severity showed that the microbial profile of severe and fatal cases clustered together. These profiles were mainly dominated by an increased relative abundance of *Parabacteroides, Lachnoclostridium*, and a reduced relative abundance of *Blautia, Faecalibacterium*, and *Ruminococcus* **(Figure 2 C)**, which were shown to be underrepresented in COVID-19[9]. There were no significant differences in the bacterial composition between COVID-19 or post COVID-19 patients. Interestingly, AC showed a higher abundance of *Coprococcus*, previously demonstrated to be associated with non COVID-19 patients[10], and *Roseburia*, which were reported to be more prevalent in healthy individuals compared to COVID-19 patients[9].

### Microbial Analysis of Saliva Samples

Alterations in the oral microbiome have previously been associated with COVID-19 and suggested as a diagnostic marker[33]. To comprehensively analyse the oro-intestinal bacterial composition, saliva samples were collected in addition to fecal samples **(Figure 1 A, Supplementary Table 6)**. In total, 160 saliva samples from 117 patients were analysed (COVID-19, N = 87, n = 117; post COVID-19, N = 13, n = 24; SC, N = 17, n = 19). Taxonomic differences on phyla level are minor with a reduced relative abundance of *Firmicutes* in COVID-19 compared to post COVID-19 and SC. Post COVID-19 showed an increased relative abundance of *Proteobacteria* and a reduction in *Actinobacteria*. Compared to SC, post COVID-19 and COVID-19 had an increased abundance of *Fusobacteria* **(Supplementary Figure 1 A)**. Overall, microbial composition between the groups showed no significant differences **(Supplementary Figure 1 B)**. Interestingly, in accordance with our findings regarding the gut bacteria, stratification of patients according to disease severity showed a significant difference in the composition of the oral microbiome (p = 0.003) **(Supplementary Figure 1 C)** as well as significant variations according to the number of complications (p = 0.001) **(Supplementary Figure 1 D)**. However, a random forest model failed to predict mortality in the setting of COVID-19-associated hospitalization for saliva samples.

### Alterations of the Gut Microbiota Correlate with Number and Type of Complications

Following the association between severity and changes in the gut microbiota, we further investigated whether microbial changes were found in terms of type and number of complications in COVID-19 and post COVID-19 patients and SC. A maximum of three complications per patient were observed. Stratifying patients according to the number of complications revealed a significant distinction between patients with no complications and patients with one or more complications, with a shift in their bacterial profile according to the number of complications (p = 0.002) **(Figure 3 A)**. Furthermore, *alpha*-diversity showed that the abundance of gut bacteria decreased with the number of complications **(Figure 3 B)**. Interestingly, *F. prausnitzii* was found to be reduced with an increased number of complications and absent in patients with three complications **(Figure 3 B)**. Consistent with the findings regarding disease severity **(Figure 2 B)**, *Parabacteroides* was increased in patients with a more complicated course **(Figure 3 B)**.

**Figure 3.**
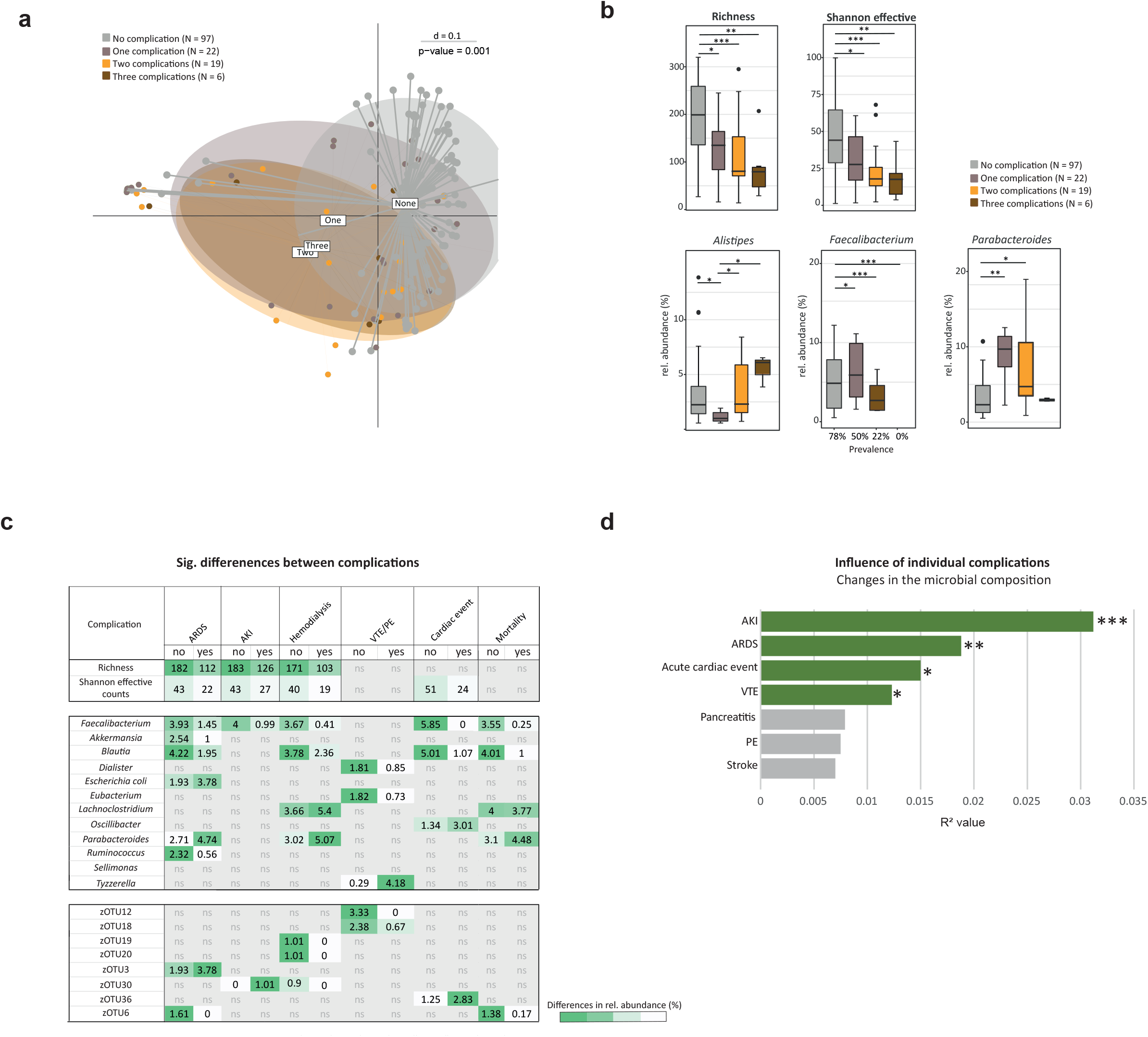
Association Between Gut Bacterial Composition and Common Complications. **A** MDS plot calculated on generalized UniFrac distance stratifying COVID-19 and post COVID-19 patients, SC and AC (T1) according to the number of complications during hospitalization. **B** Same samples as in panel A, boxplots show significant differences in *alpha*-diversity and relative abundance of taxa. *Faecalibacterium* shows differences in prevalence (p-value = 0.0002) and relative abundance (p-value ≤ 0.01), *Parabacteroides* and *Alistipes* are significantly different in their relative abundance (p-value ≤ 0.01). **C** Heatmap with taxa found to be significantly different in COVID-19, post COVID-19 and SC patients (T1) and with specific complications. Values are showing the mean relative abundance detected in patients with the complication compared to patients without complication. The color code indicates high (green) or low (white) relative abundance. **D** Multivariate permutational analysis revealed the importance of complications regarding microbial composition. Barplots are showing the R^2^ values. Green bars = significant variables (*, p ≤ 0.05, **, p ≤ 0.01; ***, p ≤ 0.001), grey = non-significant variables.

Some complications showed overlapping bacteria, which were significantly different in their relative abundance compared to patients without the corresponding complication. Patients who developed ARDS, AKI, or had hemodialysis, revealed a significantly reduced gut bacterial richness as well as Shannon effective number, which was also seen in patients with an acute cardiac event (**Figure 3 C**). Specific complications were associated with changes in the relative abundance of individual bacteria (**Figure 3 C**). Hereby, the butyrate producing *F. prausnitzii* was significantly reduced in patients with ARDS, AKI, hemodialysis, and acute cardiac events and furthermore negatively associated with mortality. *Blautia* was reduced for most complications except in patients with VTE/PE or AKI. *Parabacteroides*, on the other hand, was increased in patients with ARDS and hemodialysis and showed a positive association with mortality. Multivariate permutational analyses showed that AKI had the greatest influence on microbial changes, followed by ARDS, acute cardiac events and VTE. However, pancreatitis and stroke were not significantly contributing to microbial differences **(Figure 3 D)**.

### A Stable Gut Bacterial Composition is Correlated with A Favourable Disease Progression

During this study, 39 patients (COVID-19, post COVID-19, and SC) provided more than one stool sample, enabling the analysis of intra-individual changes during disease course. Based on generalized UniFrac distances the stability of the microbial composition of the gut was determined **(Figure 4 A)**. On average, the intra-individual distance was 0.33 ± 0.09. The microbial composition was equally dynamic between COVID-19, post-COVID-19, and SC. Compositional changes were not related with ward, nutrition, antibiotics, or disease severity. Stratifying the longitudinal data according to the number of complications supported our previous results **(Figure 3 A)** that the onset of complications during inpatient stay significantly correlated with an altered bacterial composition (p = 0.001) **(Figure 4 B)**. Even though the intra-individual distance showed no obvious grouping based on SARS-CoV-2 status, a cluster could be detected according to the simple presence or absence of complications. COVID-19 patients without any complication had a more stable microbial composition compared to patients with complications **(Figure 4 C)**. Analysis of the intra-individual microbial stability accounting for varying conditions, demonstrated the significance of environmental factors in addition to the disease state. In the context of intra-individual examination of the bacterial profiles over time, disease progression could be tracked using inflammation markers (CRP, PCT, WBC) and oxygen demand (FiO2) at the time of each stool sample. Thus, we defined a group of COVID-19 and post COVID-19 patients with severe progression. Criteria for a severe progression had to be met at least for one sampling time point (CRP ≥ 10 mg/dl, PCT ≥ 5 ng/ml, WBC ≥ 15 G/l, FiO2 ≥ 40%) and we compared this group (S-prog, N = 44) with patients not meeting these criteria (NS-prog, N = 62). Indeed, the bacterial composition of S-prog significantly differed from NS-prog **(Figure 4 D)**.

**Figure 4.**
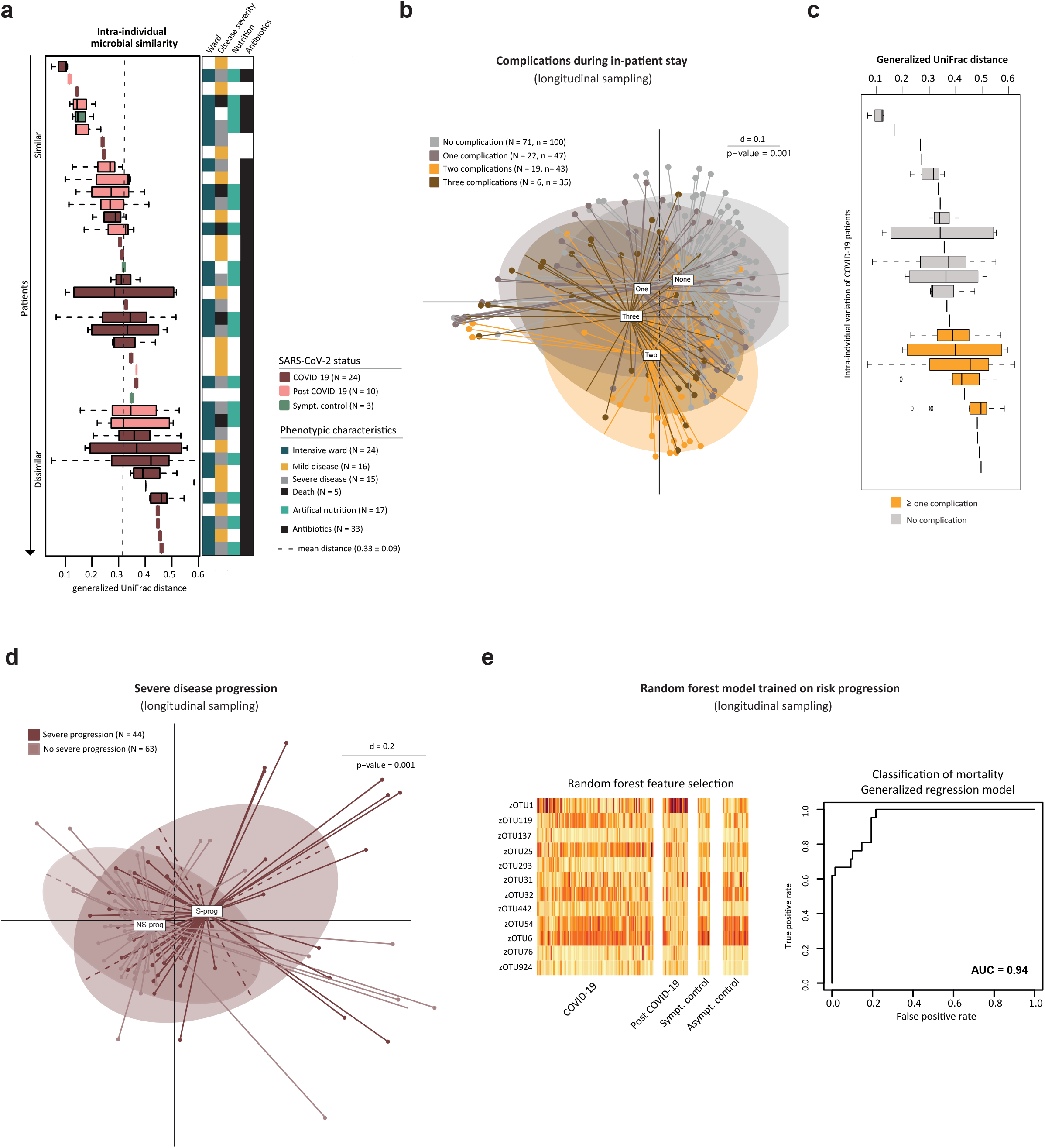
Stability of the Bacterial Composition Related to COVID-19 and Longitudinal Analysis. **A** Intra-individual generalized UniFrac distance sorted by median distance within one patient. Longitudinal sampling of at least two samples per patient with a medium of 3.5 (COVID-19), 4.6 (post COVID-19), and 2.3 (SC) samples per patient. Each box represents one patient. Dashed line shows the mean intra-individual distance over all patients (N = 39). Right color bar shows variables related to hospitalization as indicated by the legend. **B** MDS plot calculated on generalized UniFrac distances stratifying COVID-19, post COVID-19 and SC patients (all sampling time points) according to number of complications. **C** Intra-individual generalized UniFrac distances sorted by median distance within one patient of the COVID-19 cohort with a minimum of two samples (N = 25). Each box represents one patient. **D** MDS plot calculated on generalized UniFrac distance stratifying COVID-19 and post COVID-19 patients (all sampling time points) for disease progression in severe (S-prog) compared to patients not meeting these criteria (NS-prog). **E** Individual relative abundance values of random forest selected zOTUs for classification of severe disease progression grouped by SARS-CoV-2 status. ROC curve shows differentiation by mortality based on previously determined feature list.

Additionally, machine learning was applied to differentiate between S-prog and NS-prog. Towards this end, a random forest model was trained on COVID-19 patients in a 10-fold cross-validated nested approach (repeated 100 times). In total, 12 zOTUs were selected as important features: *Enterococcus durans* (zOTU1), *Streptococcus thermophilus* (zOTU119, zOTU25), *Citrobacter freundii* (zOTU137, zOTU76), *Holdemania massiliensis* (zOTU293), *Parabacteroides distasonis* (zOTU31), *D. longicatena* (zOTU32), *Lactococcus lactis* (zOTU442), *Blautia* spp. (zOTU54, zOTU6), *Lacticaseibacillus rhamnosus* (zOTU924) **(Figure 4 D, Supplementary Table 5)**. The defined signature was overlapping with zOTUs associated with disease severity within our patient population **(Figure 2 B)**. Based on this bacterial signature, a generalized linear model of all patients revealed an area under the curve (AUC) of 0.94 to predict mortality during the COVID-19 associated inpatient stay **(Figure 4 D)**. The specificity was verified by applying the signature to other outcomes, e.g. type 2 diabetes (AUC = 0.76) or presence of complications (AUC = 0.82).

## DISCUSSION

The risk of a severe disease course and complications, including thromboembolism, renal failure, and acute cardiac events is higher for COVID-19 than for influenza[34]. GI symptoms in patients with COVID-19 are associated with an increased disease severity and complications[8] and an exaggerated immune response to the virus is considered to play a crucial role in driving disease progression[15, 16]. It is well known that gut microorganisms influence the systemic immune response of their hosts through multiple crosstalk with immune cells[35, 36, 37].

In our study, we demonstrated that the bacterial composition of the gut in patients with COVID-19 disease changes with number and type of complications. Thereby, taxa known for protective and immunosuppressive properties were found to be decreased with an increasing complication rate, whereas rather pathogenic taxa were more prevalent. *F. prausnitzii*, for example, was undetectable in patients with three complications and relatively reduced in patients with AKI, hemodialysis, ARDS, cardiac event and was negatively correlated with mortality. This bacterium has anti-inflammatory properties[38, 39] and was found to have an inverse correlation with disease severity in COVID-19[9, 10]. On the other hand, the relative abundance of the genus *Alistipes* was increased with the number of complications. In terms of functionality, there is conflicting evidence to the protective or pathogenic potential of *Alistipes* in various diseases[40]. In patients with thromboembolic complications the genus *Tyzzerella* was the only significantly elevated bacterium. Interestingly, *Tyzzerella* was previously shown to be associated with an increased risk of cardiovascular diseases[41]. *Parabacteroides* was increased in patients with ARDS and hemodialysis and related to mortality. The associations of individual bacteria with the occurrence of complications suggests a potential role of the gut microbiota in the development of specific complications within COVID-19 and provide additional evidence for an involvement of the gut concerning cardiovascular risk[42] and venous thromboembolism[43, 44].

In addition, differences in the bacterial composition were found dependent on the disease severity. While the microbial profile of patients with mild diseases was comparable to controls, severe and fatal cases showed marked differences with respect to protective bacteria. Congruent with previously published studies in other countries[1, 9, 10, 45], our results confirm a link between disease severity of COVID-19 and microbiota alterations in a large German cohort. Besides an inverse correlation of *F. prausnitzii* with disease severity of COVID-19[10], *Blautia* was previously shown to be underrepresented in patients with COVID-19 and was associated with SARS-CoV-2 recovery[9]. *Fusicatenibacter* was reported to be enriched in non COVID-19 controls[45] and correlated negatively with inflammatory biomarkers in COVID-19 patients[46] and *Parabacteroides* correlated positively with disease severity[9].

To more deeply examine the associations of the gut bacteria with COVID-19 progression, we considered functional data, such as FiO2, at each time of stool collection. Thereby, the intra-individual microbial stability decreased with a higher complication rate. Based on a distinct microbial profile, the individual risk of mortality due to COVID-19 could be estimated. Thus, while disease severity, inflammatory activity, and complication rate were associated with changes in bacterial composition in COVID-19 patients, the impact of SARS-CoV-2 infection appears to be more modest, indicating that the gut plays a role in shaping severe disease progression.

Regarding the microbiota changes in the oral cavity, differences in bacterial composition related to severity and complications were observed, highlighting the importance of the bacterial oro-intestinal axis in COVID-19[33]. However, prediction of mortality was not feasible using bacterial patterns in saliva and the results were less conclusive compared to changes in the gut microbiota.

We hypothesize that changes in the microbial composition, especially of the gut, may drive disease, possibly via an involvement in the development of complications. A stable bacterial profile during hospitalization could have a favorable impact on disease progression. A healthy and diverse intestinal microbiota should, therefore, be considered in the therapeutic management of COVID-19.

Because of the often prolonged hospital stay of inpatients of 24 days on average within our cohort, multiple factors could influence the gut microbiota. These include formulated food, antibiotics, or catabolic metabolism during an ICU stay[47]. Especially in a clinically heterogeneous disease like COVID-19, these factors must be considered in the interpretation of microbiota analysis. For this reason, we carefully reviewed the results for potential confounders, including concomitant diseases and assessable factors associated with hospitalization. In this context, none of the factors examined was found to be a confounder with significant bias concerning our results. Nevertheless, patients with a severe and complicating disease, in contrast to mild cases, were mainly treated at the ICU and given antibiotics **(Supplementary Table 4)**. Thus, it cannot be ruled out that microbiota changes related to the severity and complications are also influenced by the conditions of medical treatment. It further remains unclear whether the changes in microbiota causally influenced the severity of COVID-19 and occurrence of complications, or vice versa.

Taken together, our results suggest that the gut and salivary microbiota are associated with the occurrence of individual complications in COVID-19, thereby influencing disease severity. A stable gut bacterial composition during hospitalization is associated with a more favorable clinical course. Further studies are needed to investigate direct causality between gut bacterial dysbiosis and COVID-19 and to integrate microbial patterns for prognostic and therapeutic purposes in clinical routine.

## Acknowledgments

We thank all the health care workers of Klinikum rechts der Isar as well as the CoMRI team around Christoph Spinner, MD. We are grateful to Angela Sachsenhauser, Caroline Ziegler and Lukas Mix from the Core Facility Microbiome of the ZIEL Institute for Food & Health for outstanding technical support in sample preparation and 16S rRNA gene amplicon sequencing

## Authorship Contributions

David Schult, MD (Conceptualization: Lead; Project administration: Lead; Supervision: Lead; Investigation: Equal; Methodology: Lead; Visualization: Supporting; Validation: Equal; Manuscript – writing: Lead, Manuscript – review & editing: Lead), Sandra Reitmeier, PhD (Conceptualization: Equal; Project administration: Equal; Investigation: Equal; Methodology: Lead; Visualization: Lead; Validation: Equal; Manuscript – writing: Lead, Manuscript – review & editing: Lead; Formal analysis: Lead), Plamena Koyumdzhieva, cand. med. (Conceptualization: Equal; Investigation: Lead; Methodology: Supporting; Visualization: Supporting; Validation: Equal; Manuscript – writing: Equal, Manuscript – review & editing: Supporting), Tobias Lahmer, MD, Johanna Erber, MD, Marina Frolova, Julia Horstmann, Lisa Fricke, Juliane Kager, MD, Katja Steiger, MD, Moritz Jesinghaus, MD (Investigation: Supporting), Moritz Middelhoff, MD, Jochen Schneider, MD (Conceptualization: Supporting), Klaus-Peter Janssen, PhD (Resources: Supporting; Manuscript – review & editing: Supporting), Ulrike Protzer, MD (Resources: Supporting), Klaus Neuhaus, PhD (Resources: Equal; Validation: Supporting; Manuscript – review & editing: Equal), Roland M. Schmid, MD (Resources: Equal; Funding acquisition: Equal), Dirk Haller, PhD (Conceptualization: Supporting, Resources: Equal; Validation: Supporting; Funding acquisition: Equal; Manuscript – review & editing: Equal), Michael Quante, MD (Conceptualization: Lead; Project administration: Equal; Supervision: Lead; Funding acquisition: Lead; Manuscript – review & editing: Lead).

## Data Availability and Data Transparency Statement

FASTQ files of the 16S rRNA gene sequencing is available under SRA accession number PRJNA756849 (https://www.ncbi.nlm.nih.gov/bioproject/PRJNA756849/).

## Abbreviations

AB T1: Antibiotic therapy at the time of the first stool sampling
AC: Asymptomatic controls
AKI: Acute kidney injury
AP: Alkaline phosphatase [U/l]
ARDS: Acute respiratory distress syndrome
Asympt.: Asymptomatic
COVID-19: Corona virus disease 2019
CRP: C-reactive protein [mg/dl]
FiO2: Fraction of inspired oxygen (%)
GGT: Gamma-Glutamyltransferase [U/l]
GI: Gastrointestinal
Hb: Hemoglobin [g/dl]
i.v.: intravenous
IBD: Inflammatory bowel disease
ICU: Intensive care unit
ICU all T: Intensive care stay regarding all time points of stool sampling
ICU T1: Intensive care stay at the time of the first stool sampling
N: Number of patients
n: Number of samples
NA: not available
PA: Pressure assisted
PC: Pressure controlled
PCT: Procalcitonin [ng/ml]
PE: Pulmonary embolism
Rel.: Relative
S.p.: Status post
SARS-CoV-2: Severe acute respiratory syndrome coronavirus 2
SC: Symptomatic pneumonia controls
Sig.: Significant
Sympt.: Symptomatic
T1: First sampling time point
T2D: Type 2 diabetes mellitus
Trp T: High-sensitive troponin T [ng/ml]
TS: Tracheostomy
VTE: Venous thromboembolism
WBC: White blood cells counts [G/l]
zOTU: Zero-radiation operational-taxonomic units

## Figure and Table Legends

**Supplementary Figure 1 Microbial Profile of Sputum Samples**

**A** Upper plot shows the taxonomic distribution based on phyla level over all patients (T1). Stacked barplots represent phyla composition stratified by SARS-CoV-2 status. **B-D** MDS plot calculated on generalized UniFrac distance stratifying at T1 according to **B** SARS-CoV-2 status, **C** disease severity, **D** number of complications during hospitalization.

**Supplementary Table 1 Classification of the Antibiotic Therapy given to Patients during their Inpatient Stay**

**Supplementary Table 2 Results of Confounding Analysis**

**Supplementary Table 3 Distribution of Factors with Possible Impact on the Gut Microbiota**

**Supplementary Table 4 Antibiotics and Intensive Care According to COVID-19 Severity and Complications**

**Supplementary Table 5 Taxonomic Classification of zOTUs in Fecal Samples**

**Supplementary Table 6 Taxonomic Classification of zOTUs in Saliva Samples**

