## Supplementary figures and images for "Gut Bacterial Dysbiosis and Instability is Associated with the Onset of Complications and Mortality in COVID-19"

### Supplementary Figure 1

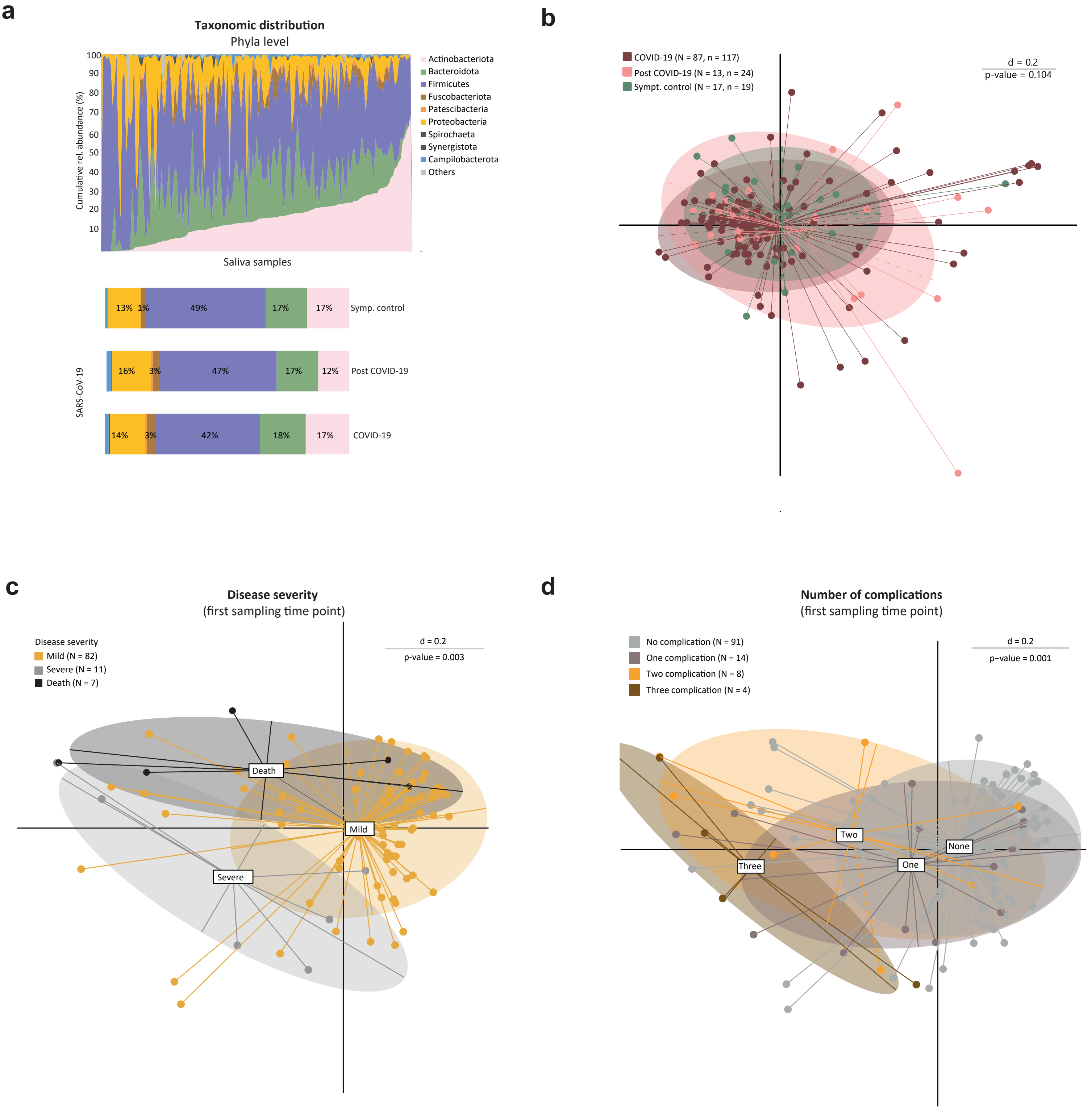
