## Supplementary Table 1 for "Gut Bacterial Dysbiosis and Instability is Associated with the Onset of Complications and Mortality in COVID-19"

| **Group** | **Antibiotics** | **Spectrum of activity** | **Special features** |
| --- | --- | --- | --- |
| **1 = Broad-spectrum** | **- Beta-lactam antibiotics** (Acylaminopenicillins, Aminopenicillins, Carbapenems)  **- Tetracyclins**  **- Levofloxacin**  **- Moxifloxacin** | - Anaerobic bacteria  - Gram-positive bacteria  - Gram-negative bacteria |  |
| **2 = Narrow-spectrum** | - **Linezolid**  **- Daptomycin**  **- Flucloxacillin**  **- i.v. Vancomycin** | - Gram-positive bacteria |  |
| **3 = Cephalosporins** | **- Cephalosporins** | - Anaerobic bacteria  (partly)  - Gram-positive bacteria  - Gram-negative bacteria | - lack of activity against enterococci |
| **4 = Others** | **- Gentamicin**  **- Ciprofloxacin**  **- Azithromycin** | - Gram-positive bacteria (partly for Aminoglycosides),  - Gram-negative bacteria |  |
